# Mutational hotspots lead to robust but suboptimal adaptive outcomes in certain environments

**DOI:** 10.1101/2023.06.07.543998

**Authors:** Louise M. Flanagan, James S. Horton, Tiffany B. Taylor

## Abstract

The observed mutational spectrum of adaptive outcomes can be constrained by many factors. For example, mutational biases can narrow the observed spectrum by increasing the rate of mutation at isolated sites in the genome. In contrast, complex environments can shift the observed spectrum by defining fitness consequences of mutational routes. We investigate the impact of different nutrient environments on the evolution of motility in the bacteria *Pseudomonas fluorescens* Pf0-2x (an engineered non-motile derivative of Pf0-1) in the presence and absence of a strong mutational hotspot. Previous work has shown that this mutational hotspot can be built and broken via six silent mutations, which provide rapid access to a mutation that rescues swimming motility and confers the strongest swimming phenotype in specific environments. Here, we evolved a hotspot and non-hotspot variant strain of Pf0-2x for motility under nutrient rich (LB) and nutrient limiting (M9) environmental conditions. We observed the hotspot strain consistently evolved faster across all environmental conditions and its mutational spectrum was robust to environmental differences. However, the non-hotspot strain had a distinct mutational spectrum that changed depending on the nutrient environment. Interestingly, while alternative adaptive mutations in nutrient-rich environments were equal to, or less effective than, the hotspot mutation, the majority of these mutations in nutrient-limited conditions produced superior swimmers. Our competition experiments mirrored these findings, underscoring the role of environment in defining both the mutational spectrum and the associated phenotype strength. This indicates that while mutational hotspots working in concert with natural selection can speed up access to robust adaptive mutations (which can provide a competitive advantage in evolving populations) they can limit exploration of the mutational landscape, restricting access to potentially stronger phenotypes in specific environments.

**DATA SUMMARY:** All data is available at OSF: DOI 10.17605/OSF.IO/8BT2W, and in supplementary materials.

## INTRODUCTION

Mutational hotspots - regions of the genome that mutate at a higher frequency than expected by chance alone - have been reported widely in nature, especially in relation to phenotypic changes (1–5). They are responsible for many parallel and convergent evolutionary events such as evolution of rifampicin resistance via mutations in the same RNA polymerase subunit across bacteria (6) to repeated instances of cancer-causing mutations in humans (3,7). When a strong mutational hotspot is found at an adaptive genomic position where mutation will bolster a phenotype under selection, the outcome can be remarkably predictable evolution (8).

Understanding the effect of mutational hotspots’ presence or absence on adaptive outcomes has not yet been widely studied. What are the consequences of having a mutational hotspot or not in a system where selection will predictably converge on the same phenotype? Will absence of a hotspot reveal a less constrained mutational landscape that lends itself to discovery of fitter mutants, or will the presence of a hotspot provide its own advantages through rapid evolution and predictable adaptive outcomes? Mutational hotspots elevate the mutation rate at a particular location in the genome and the potency of these hotspots can vary due to multiple interacting molecular features (9).

In recent years, much attention has been given to the principle of predicting evolution, and how this could be used to gain a greater understanding of future adaptive phenotypes (10–15). Current global emergencies, such as pandemics and rapid climate change, highlight an immediate application for such knowledge, with clear advantages gained from being able to anticipate how populations will adapt. However, we must first expand our understanding of the role the environment plays in defining predictable outcomes. Many experimental evolution studies focus on predictability under singular, defined conditions (16–19). However, other studies have highlighted the importance of including varied environments in experimental evolution, as it has an undeniable influence on the evolutionary trajectory and the subsequent fitness of mutants in alternative environmental conditions (20–24).

In this work, we utilise a model system that allows direct empirical tests into the interaction of varying environments with mutational biases. Here we look at the consequences of having a mutational hotspot and not having one in varying environments by taking two nearly identical strains of bacteria, one with an engineered mutational hotspot and one without, and putting them under strong selective pressure to restore the same lost motility phenotype. Through this we can see how the presence or absence of the hotspot, along with differing nutrient regimes, determines several outcomes: ease of access to an adaptive phenotype, the quality of that adaptive phenotype, and how these factors interact to influence competition between the hotspot and non-hotspot strains. We also investigate the interaction of nutrient environment with these varying mutational biases.

In previous work, Horton et al (8) showed that a mutational hotspot can be built or broken with just a few synonymous mutations. The immotile derivatives of two strains of *Pseudomonas fluorescens*: SBW25 (which naturally contains a mutational hotspot in *ntrB*) and Pf0-1 (no native mutational hotspot in *ntrB*), were used to test this. These strains were genetically engineered to lose their swimming phenotype through deletion or disruption of the master regulator gene for the flagellum, *fleQ*. They reliably restored motility after just a few days of selection through starvation on 0.25% swimming agar plates (25,26). They rescued motility through co-option of a distantly related master regulator NtrC, from the nitrogen regulatory (*ntr*) network to partially take over the function of the missing FleQ. This co-option initially results from hyperactivation of NtrC achieved via mutations within other genes in the nitrogen regulatory pathway. In the strains with a mutational hotspot, Horton et al. (8) frequently observed an A to C transversion at nucleotide 289 in the gene *ntrB* (*ntrB* A289C). In strains without a hotspot, mutations occurred in genes *ntrB, glnA,* and *glnK*, all genes within the *ntr* pathway. In an extension of this work, Horton et al. (27) were able to show that this hotspot could facilitate rapid and predictable access to the strongest observed motility phenotype in a nutrient rich environment. However, we know that mutations in the *ntr* pathway have pleiotropic consequences for how the variants can grow in different nitrogen environments (25). The interplay between these two factors in defining adaptive outcomes is yet to be explored.

In this work we take derivatives of the Pf0-1 strain, Pf0-2x (an engineered non-motile variant without a native mutational hotspot in the *ntrB* gene) and Pf0-2x-sm (an engineered non-motile variant with an engineered hotspot in the *ntrB* gene) to ascertain the effect of having a hotspot or not in an experimentally evolved system. These strains are genetically identical aside from 6 synonymous mutations in the *ntrB* locus which confers a hotspot in the Pf0-2x-sm strain. Pf0-2x-sm was chosen for this study instead of the naturally occurring hotspot strain SBW25 in order to remove the variable of genetic background. Genetic background has been repeatedly reported to lead to confounding and inconsistent results (28,29). In this study we aimed to uncover the effects of different nutrient regimes on the evolution of motility rescue in the presence or absence of a mutational hotspot.

## METHODS

### Strains and Culture Conditions

All *Pseudomonas fluorescens* strains and derivatives (Pf0-2x, Pf0-2x-sm, Pf0-1 *ΔfleQ*, Pf0-2x-sm *ntrB* A289C *glmS*::Kan^R^) were grown at 27°C for approximately 48h on solid LB (lysogeny broth) agar supplemented with either ampicillin (100 µg/µl), streptomycin (50 or 250 µg/µl), or kanamycin (50 µg/µl) before use in downstream experiments (Table S.1).

Overnight cultures were grown at 27°C at 180 rpm in 10 ml LB or M9 minimal media (containing glucose and 7.5 mM NH_4_) supplemented with 10 µl of the appropriate antibiotic.

### Motility Evolution Assays

A single colony of immotile *P. fluorescens* was inoculated into 0.25% LB and M9 agar plates (except for 3 Pf0-2x mutant lines where a 1 µl stab of OD_595nm_ 1 cells was used. See Supplementary Materials for details). Preparation of these plates has been outlined previously (8). Plates were monitored daily to check for the restoration of a motility phenotype. Once motility was restored, cells from the outermost edge of the motile zone were sampled and streaked onto solid agar supplemented with the appropriate antibiotic and incubated as described above. A single colony from these plates was then restreaked onto another solid agar plate to ensure genetic uniformity and incubated again. A single colony from this plate was then used to make an overnight culture and cryopreserved in 20% glycerol at -80°C.

### Sanger Sequencing of Mutant Lines

To identify the mutations that restored motility, DNA of candidate genes (*ntrB*, *glnA*, *glnk*, as identified in Taylor et al. (25) and Horton et al. (8) from each evolved line were amplified via colony PCR (the list of primers used is available in Table S.2). PCR products were run on a 1% agarose gel to assess amplification, and successful PCR products were cleaned up using Monarch® PCR and DNA Clean-Up Kit (New England Biolabs) as per kit instructions. Samples were sent for Sanger sequencing via Eurofins Genomics and were prepared as per company instructions (5-10ng/µl of cleaned up DNA product plus 2 µl of primer to a total volume of 17 µl).

### Whole Genome Sequencing of Mutant Lines

In instances where the mutations that restored motility could not be found through targeted Sanger sequencing, samples were sent for whole genome sequencing (WGS). Genomic DNA was extracted using the ThermoFisher GeneJET Kit as per kit instructions with the following amendments to increase yield for Pf0-2x lines: 1 ml of neat overnight culture was spun at 5000 xg for 10 min. The cell pellet was washed and resuspended in PBS and then spun again. The 56°C incubation step was extended to an hour and vortex times were doubled to what was instructed. Genomic DNA integrity was checked, and concentration was measured through Qubit and Nanodrop spectrophotometry as well as through electrophoresis of a 1% agarose gel with the prepared samples. Prepared gDNA was sent for Illumina sequencing to Microbes NG (Birmingham, UK) and SeqCenter (Pittsburgh, USA) with a minimum of 30x coverage. Illumina paired end read data of the evolved Pf0-2x lines was aligned to the *P. fluorescens* Pf0-1 reference genome (30) using Integrated Genome Viewer (31). This was used to scan for large deletions or insertions. Presence of SNP mutations was analysed using the variant calling software SNIPPY using default parameters (32).

### Swimming and Growth Assays of Evolved and Unevolved Mutant Lines

To assess the swimming speed and metabolic profile of the evolved and ancestral *P. fluorescens* lines, swimming race assays and growth curve assays were performed respectively. Cultures of each line were grown overnight in LB. Cells were diluted with PBS to an OD_595nm_ of 1. For swimming race assays 1 µl of these diluted cells were used to inoculate the centre of a 0.25% agar plate, supplemented with the appropriate nutrient media. The pipette tip was inserted into approximately half the depth of the agar before allowing the cells to effuse. This was replicated at least 3 times for each evolved line in both LB and M9. These plates were incubated at 27°C for 24 h. Photos were taken of the plates and measurements of the distance swam were taken from these photos. The surface area was calculated using the following equation: *A* = π*r*^2^, and was square root transformed.

For growth curve assays 1 µl of diluted culture was added to 99 µl of LB or M9 broth in a Corning 96 well flat-bottomed transparent plate. These were run for 24 h, at 180 rpm and 27°C in a spectrophotometric plate reader (TECAN or Multiskan SkanIt FC), with readings every 10 minutes or 1 h respectively. Area under the curve was calculated to measure growth profile using the R package *growthcurver* (33).

### Competition Assays

Evolved hotspot and non-hotspot mutants were competed against each other to determine which have the strongest swimming phenotypes and metabolic profiles. To allow for distinction between the hotspot and non-hostpot lines, Pf0-2x-sm *ntrB* A289C was marked with a kanamycin resistance cassette in the *glmS* locus (Pf0-2x-sm *ntrB* A289C *glmS*::Kan^R^) using the mini Tn7 transposition system as outlined by Choi et al. (34) and Liu et al. (35). Overnight cultures of the hotspot mutant Pf0-2x-sm and the non-hotspot mutants Pf0-2x A141P *ntrB* and ΔLVRG *ntrB* were grown in LB and OD_595nm_ adjusted to a value of 1 using PBS. For the competitive swimming assays, 0.5 µl of the hotspot mutant and 0.5 µl of either non-hotspot mutant was co-inoculated into the centre of a 0.25% agar plate. This was replicated 6 times for each in both LB and M9. Plates were incubated at 27°C until the agar became saturated with swimming bacterial cells (>7 days). The agar for each plate was then disrupted using a sterile spreader and 25 ml pipette, using PBS to help liquify the agar. This solution was then transferred to 50 ml falcon tubes and topped up to equal volumes with PBS. These cells were serially diluted with PBS and spotted (20 µl in triplicate) onto selective agar (LB streptomycin (250 µg/µl), and LB kanamycin (50 µg/µl) plus streptomycin (250 µg/µl)). Pf0-2x-sm *ntrB* A289C *glmS*::Kan^R^ only grows on LB kanamycin and streptomycin plates so can be distinguished from Pf0-2x A141P *ntrB* and ΔLVRG *ntrB*. Colony forming units were counted to determine overall abundance of each line on the entire plate (Fig.S.1). For the growth competitions to measure metabolic fitness, overnight cultures of the hotspot and non-hotspot mutants were inoculated 50:50 into 10 ml of either LB or M9, giving a starting OD_595n_ of 0.01, and incubated at 27°C 180 rpm for 24 h.

### Competitive Evolution Assays

Immotile hotspot (Pf0-2x-sm) and non-hotspot (Pf0-1 *ΔfleQ*) lines were competed against each other to determine which restored motility first. Pf0-1 *ΔfleQ* was used here instead of Pf0-2x so that cells could be differentiated post-competition (Table S.1). Cells from overnight cultures of each competition line were OD_595nm_ adjusted to 1 and co-inoculated in 50:50 ratio (0.5 µl : 0.5 µl) into 0.25% agar plates and incubated as before. Plates were monitored daily to check for restoration of motility. When evolution of motility was observed samples of cells from the leading edge of the motile zone were taken and streaked onto selective agar to determine which bacterial line(s) were present (Fig.S.2).

### Data Analysis

All statistical tests were performed in R and all plots were made using the *ggplot2* package (36). Datasets were tested for normality through data visualisation (histograms and qqplots). Data that were not deemed normal initially were analysed using non-parametric tests, or log transformed to achieve a normal distribution. To compare medians for two or more groups, a Kruskal Wallis test was used. To compare means of two of more groups an ANOVA was performed. Fisher’s Exact test was used to compare differences between categorical variables when the sample size was small (one of the expected outcomes was predicted to be <5). Pearson’s Chi Squared test was used to compare categorical variables when the sample size was larger (expected outcomes for each variable >5). Contingency tables were generated within the default parameters for Fishers Exact and Pearson’s Chi Squared test in R and were checked before use. *p*<0.05 was taken to indicate a significant result for all tests.

## RESULTS

### The presence of the mutational hotspot increases the rate of evolutionary rescue of the motility phenotype in immotile *Pseudomonas fluorescens* strains

This work utilised two genetically identical immotile strains, Pf0-2x and Pf0-2x-sm. The only difference between these strains is six synonymous mutations in the *ntrB* locus (G276C, T279C, G285C, G291C, G294T and C300G) that define a mutational hotspot (a genetic region with an elevated mutation rate) (8). They are metabolically equivalent with no significant difference in growth rate in the nutrient environments used in this study (LB, *p*= 0.4433, M9 *p*= 0.4433, Kruskal Wallis Rank Sum Test, see Fig.S.3).

To assess the effect of the mutational hotspot on the rate of evolutionary rescue of the motility, we placed immotile bacteria under selection to restore motility through starvation on 0.25% swimming agar plates supplemented with either LB or M9. Both lines of bacteria re-evolve motility after a few days, but the speed of recovery of this phenotype varies between them (Fig. 1). The hotspot strain, Pf0-2x-sm, restores motility on LB and M9 within 2-6 days, whereas Pf0-2x takes significantly longer across both nutrient environments: 3-9 days in LB, and 3-13 days in M9 (*p*< 0.001, Kruskal Wallis Rank Sum Test).

**Fig. 1.**
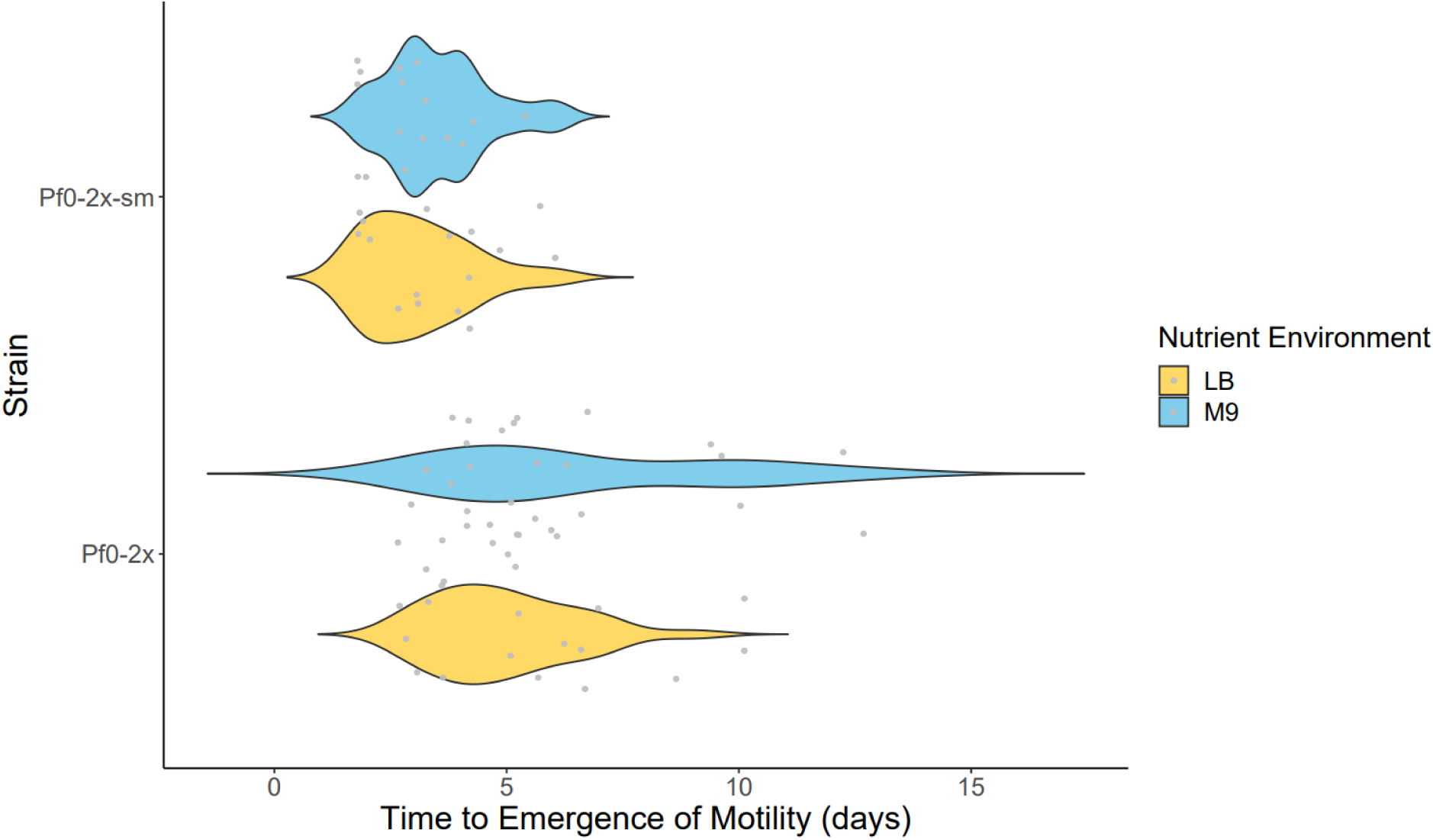
Time taken to restore the motility phenotype in the hotspot (Pf0-2x-sm) and non-hotspot (Pf0-2x) lines in complex (LB) and defined (M9) nutrient environments. Violin plots showing time taken for Pf0-2x-sm *(n* = 31) and Pf0-2x *(n* = 51) (immotile derivatives of *Pseudomonas fluorescens* strain Pf0-1) to evolve motility on 0.25% motility agar across different nutrient environments, LB (yellow) and M9 (blue).

Interestingly, the presence of the mutational hotspot (Pf0-2x-sm) does not affect the speed of motility re-evolution across different nutrient environments (*p*= 0.2385, Kruskal Wallis Rank Sum Test). However, in the absence of the hotspot (Pf0-2x) motility is restored at significantly different rates across LB and M9 (*p*= 0.03, ANOVA). These results suggest that the presence of the mutational hotspot in Pf0-2x-sm can significantly decrease the time taken for a motile phenotype to evolve. But does this hotspot offer a competitive advantage across different nutrient environments?

### Nutrient environment determines mutational spectra in the nitrogen regulatory pathway in the absence, but not the presence, of the hotspot

The presence of the mutational hotspot confers parallel evolution of motility-restoring mutations – with an A to C transversion at site 289 in *ntrB* (hereafter *ntrB* A289C) overwhelmingly dominating evolved motile lines (8). In this study, we further investigated the impact of nutrient environment on the mutational spectra in both the hotspot and non-hotspot strains (data included from Horton et al. (37)). The common *ntrB* A289C mutation has different fitness consequences in different nitrogen environments, with a notable reduced doubling time where ammonium is the sole nitrogen source (25), due to its impact on nitrogen assimilation. Here, we placed Pf0-2x and Pf0-2x-sm under directional selection for motility in complex (LB) and defined (M9) nutrient environments. LB is a rich nutrient media with multiple sources of nitrogen, whereas M9 only has ammonium available to utilise. Targeted loci *ntrB*, *glnA* and *glnK*, known as likely candidate genes for mutation from previous work (8,25) were sequenced using Sanger sequencing (Eurofins). Whole genome sequencing (MicrobesNG, SeqCenter) was performed on any lines where mutations in the candidate genes were absent and to check for any other background mutations that may also be influencing the phenotype. None of note were found, though the motility restoring mutations for some lines remain unidentified.

As seen previously, in the mutational hotspot line (Pf0-2x-sm) we see parallel evolution in motility restoring mutations at nucleotide 289 in *ntrB* (8). Within the hotspot line we also see other rare mutations in a comparatively small proportion of replicates (Fig. 2 (a)). All mutations were in *ntrB* and rare mutations include an in frame-deletion (Δ216-230), and a G328A substitution, both in LB, plus one unidentified mutation in M9. In the non-hotspot strain (Pf0-2x), we observe a wide range of mutations that can restore motility. These mutations occur across the nitrogen regulatory pathway genes *ntrB*, *glnA*, and *glnK* and vary between SNPs and indels in different domains of the protein products of these genes.

**Fig. 2.**
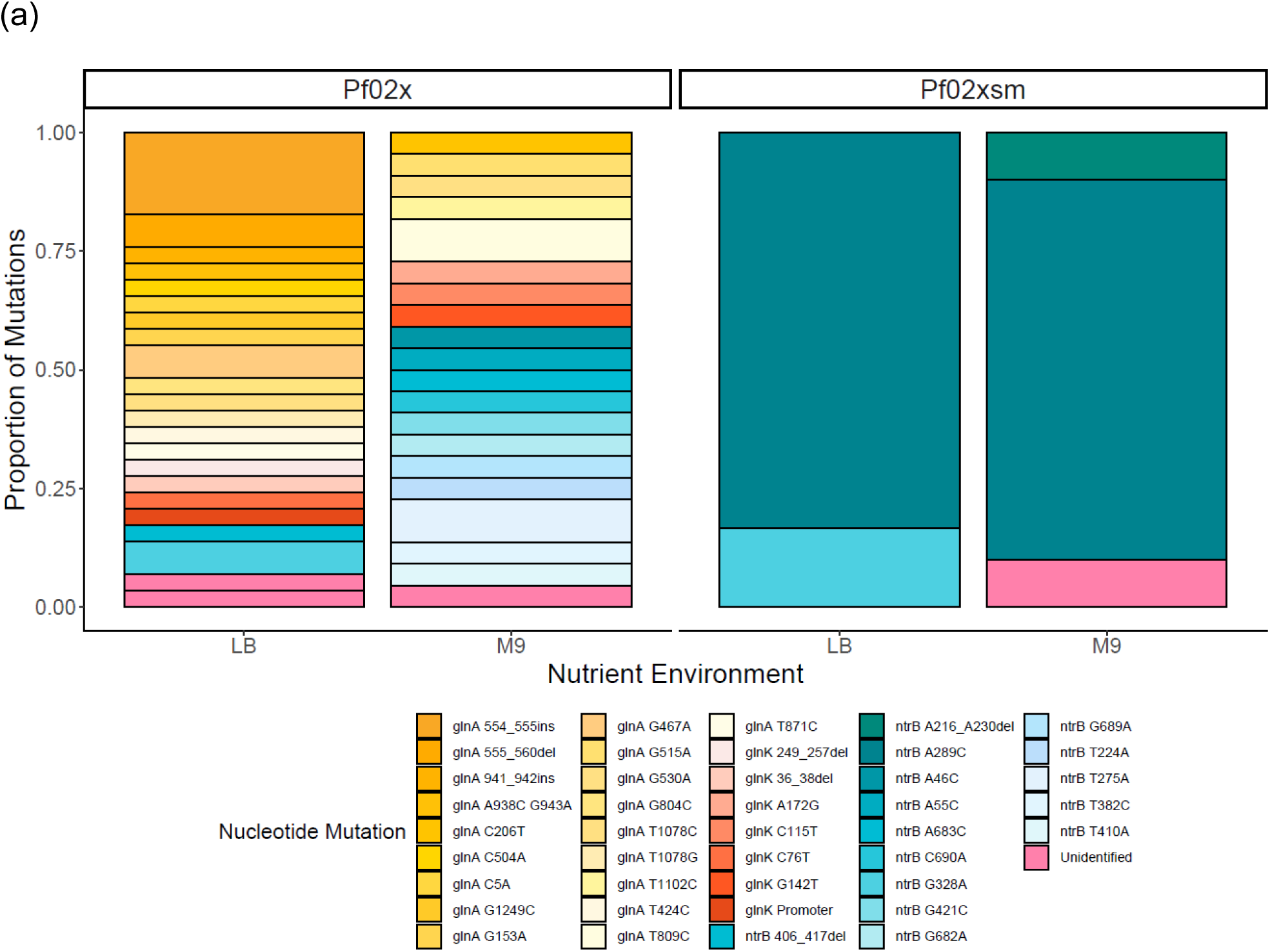
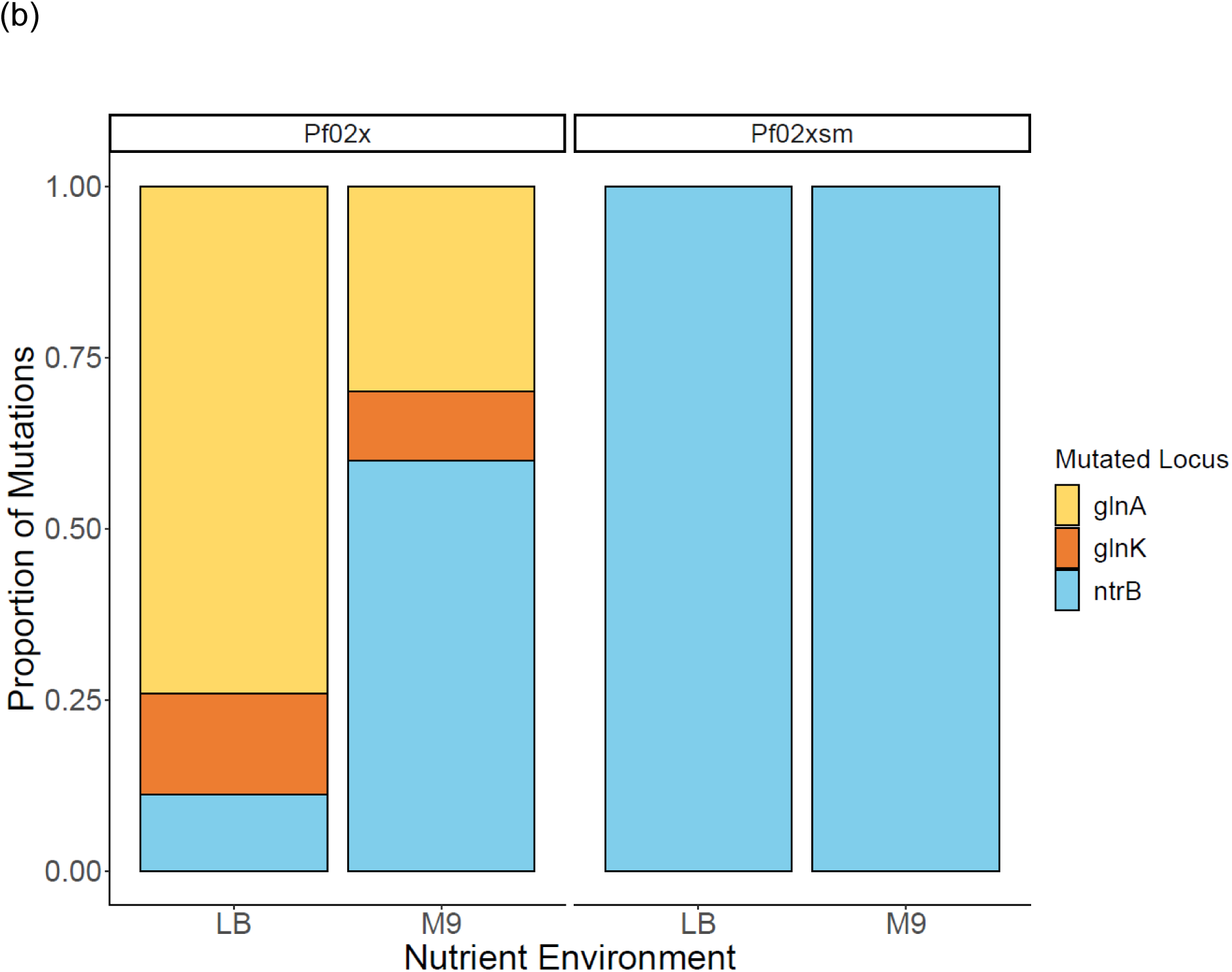
Motility restoring mutations in Pf0-2x-sm (hotspot strain) and Pf0-2x (non-hotspot strain) across different nutrient environments. (a) The proportion of each mutation conferring motility rescue across Pf0-2x and Pf0-2x-sm is shown on the y-axis, with each different observed mutation corresponding to a different colour. We find strong parallel evolution at site 289 in gene *ntrB* in the presence of the hotspot (Pf0-2x-sm). In the absence of the hotspot (Pf0-2x), we see a much wider mutational spectrum that covers multiple genes in the nitrogen regulation pathway (*ntrB*, *glnA* and *glnK*). Pf0-2x-sm has a strong mutational hotspot at the A289C site in the *ntrB* locus to restore motility in LB (*n*=6) and M9 (*n*=10). In Pf0-2x there is a much wider range of mutations that can restore motility across both LB (*n*=29) and M9 (*n*=21) nutrient environments. (b) Only mutations in *ntrB* are observed in Pf0-2x-sm. In the non-hotspot line, Pf0-2x, we see mutations in *ntrB, glnA,* and *glnK*. There is a bias towards mutations in *glnA* in LB (20/27) and a bias towards *ntrB* mutations in M9 (12/21). (Data included here is from Horton et al. (37)).

Notably, in the hotspot strain, all mutations were in *ntrB* across both LB and M9, whereas we see a bias towards mutations in certain loci dependent on the nutrient environment in which they were evolved in the non-hotspot line (Pf0-2x) (Fig. 2 (b)). In the complex nutrient environment (LB) we see a bias towards mutations in *glnA* (20/27) Whereas in the defined media environment (M9) a large proportion of *ntrB* mutations are observed (12/21). Overall, we find a significant difference in the mutation spectra between the LB and M9 nutrient environment in the non-hotspot strain (Pf0-2x) (*p*=0.002, Pearson’s Chi Squared Test), and also between both non-hotspot and hotspot (Pf0-2x-sm) strains across both LB and M9 nutrient environments (*p*=0.00002, Pearson’s Chi Squared Test). These results suggest that the nutrient environment can significantly influence the mutational spectra of motility rescuing mutations via the nitrogen regulatory pathway genes in *P. fluorescens*, however, presence of the mutational hotspot can override this trend and promote parallel evolution of motility-restoring mutations at nucleotide 289 independent of the nutrient environment.

### Motile lines evolved from non-hotspot lines show greater variability in swimming strength and can access mutations that confer a stronger phenotype in the nutrient limited environment

The hotspot mutant Pf0-2x-sm has been shown to restore the motility phenotype faster than Pf0-2x in both LB and M9 media. However, we aimed to determine if different mutations in the nitrogen regulation pathway would confer varying strengths of swimming phenotype. To test this, we inoculated all evolved motile mutants derived from Pf0-2x and the Pf0-2x-sm hotspot mutant in 0.25% agar (LB & M9) with cells normalised to an OD _595nm_ of 1 from an overnight culture, and measured swimming distance in 24 h (Fig. 3). Our overarching question was whether the common hotspot mutation (*ntrB* A289C) conferred the fittest phenotype, and as such the distance swam by all mutants is plotted relative to swimming speed of Pf0-2x-sm *ntrB* A289C (Fig. 3). In both the LB and M9 swimming assays, the distance swam by the non-hotspot line was significantly different to the hotspot line (LB *p*<0.0001; M9 *p*<0.0001 Kruskal Wallis Test).

**Fig. 3.**
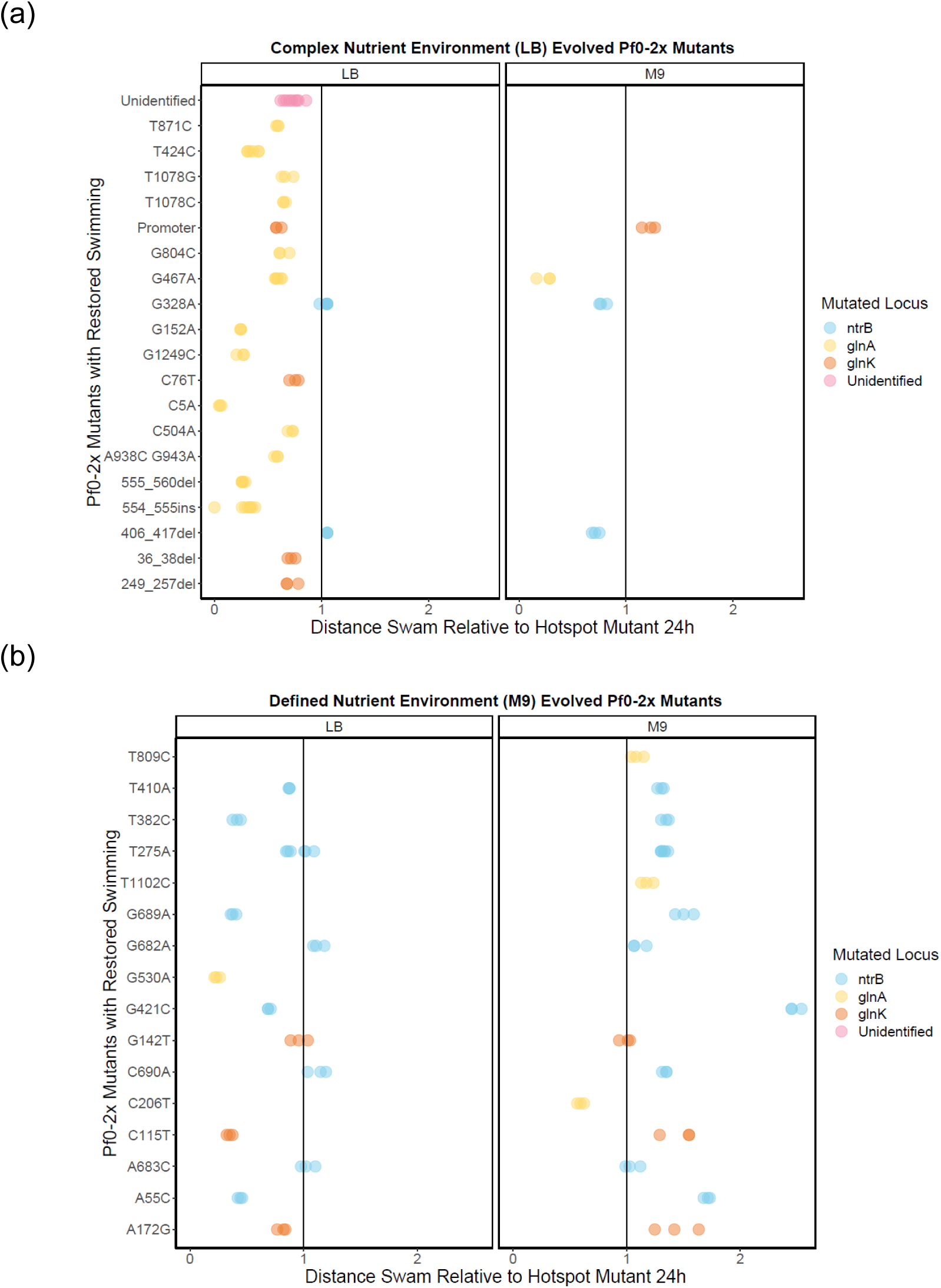
The strength of restored swimming phenotypes evolved from the non-hotspot strain, relative to the common hotspot mutant, in LB and M9 nutrient environments. The swimming distance of Pf0-2x evolved mutants are plotted relative to the Pf0-2x-sm *ntrB* A289C hotspot mutant (indicated by the black line). Data points represent Pf0-2x evolved swimmers and are distinguished by colour based on the locus where the mutation to restore motility occurred. All replicates (n≥3) are plotted separately. (a) LB-evolved Pf0-2x mutants were assayed for their strength of swimming phenotype in their adapted environment, LB, as well as the minimal nutrient environment, M9. (b) The swimming phenotype of the M9-evolved Pf0-2x mutants was assayed in both the adapted M9 environment and alternate LB nutrient environment.

To assess whether the mutants displayed differences in swimming phenotype strength in the environment in which they were evolved, we tested them in their adapted environment and in the alternate nutrient environment. LB-evolved Pf0-2x mutants swam approximately the same or worse than the hotspot mutant (Fig.3 (a)), with only *ntrB* G329A and *ntrB* Δ407-418 mutants assayed in LB swimming faster than the common hotspot mutation (*ntrB* A289C). We subsequently assessed the mutants’ swimming phenotypes in the alternate environment to look for environment-specific effects. We observed that many of the LB-evolved mutants did not swim in the M9 nutrient environment (17/21), which were predominantly *glnA* mutants (12/17). Non-motile M9-evolved *glnA* mutants were also observed when assayed in LB (2/3 *glnA* mutants were immotile in LB 0.25% agar).

Among the LB evolved mutants assayed in M9, only one mutant (featuring a mutation in the promoter region of *glnK*) swam faster than the hotspot mutant. Among the M9-evolved Pf0-2x mutants, 3/13 swam faster than the hotspot mutant in LB (Fig. 3 (b)). In the M9 assay, 13/16 swam faster than the common *ntrB* A289C hotspot mutant. The fastest non-hotspot mutant across both LB and M9 evolved lines was *ntrB* G421C, assayed in its adapted environment, M9. These results demonstrate that the motile mutants evolved from non-hotspot lines (Pf0-2x) exhibit greater variability in the strength of the swimming phenotype and can access mutations that confer stronger swimming phenotypes.

### The hotspot genotype outcompetes the non-hotspot genotype in the race to evolve motility when sharing the same ecological niche

The hotspot line, Pf0-2x-sm, evolves the motility phenotype faster than the non-hotspot line Pf0-2x. Yet the non-hotspot line can access mutations that confer a stronger swimming phenotype. This raises the question as to which factor, rate of evolutionary rescue or breadth of mutational spectrum, is most important in defining competitive fitness in an evolving population. To address this, we co-inoculated Pf0-1Δ*fleQ* and Pf0-2x-sm at equal cell densities, in both LB (*n*=6) and M9 (*n*=6) 0.25% agar and incubated them at 27°C until they evolved motility. Pf0-1*ΔfleQ* was used in this assay instead of Pf0-2x to allow post competition differentiation. Pf0-2x has a transposon disruption in *fleQ*, knocking-out function, whereas Pf0-1*ΔfleQ* has a gene deletion (Table S.1.), however they are functionally equivalent in terms of growth rate (LB *p*=0.4433, M9 *p*=0.4433, Kruskal Wallis Rank Sum Test, Fig.S.4). Once a motile zone emerged on the plate, a sample was taken and streaked onto selective agar (streptomycin 250 µg/µl to select for Pf0-2x-sm) to determine which bacterial line was present at the leading edge of the motile colony.

In the LB environment, the hotspot strain (Pf0-2x-sm) was the only strain present at the leading edge of the motile zone in every instance (Fig.S.6a). In the M9 environment, the hotspot strain was present at the leading edge in five of the six replicates but in one replicate, the non-hotspot line evolved at the same time on the same plate, giving rise to two separate motile zones (Fig.S.6b). All the evolved motile mutants sampled from the leading edge had their *ntrB* locus sequenced, and in all replicates, the *ntrB* A289C mutation was identified as the mutation conferring motility. These results suggest that in this context, the rate of evolutionary rescue is the most important factor defining competitive fitness when evolving in the same ecological niche. The ability of the non-hotspot strain to access mutations that confer a stronger wiming phenotype may result in an advantage between evolved populations. However, when evolving in the same environment as the hotspot line, its slower rate of evolution puts it at a disadvantage.

### The fastest evolved non-hotspot swimmers persist and outcompete the common hotspot mutant in competitive swimming assays

While presence of the hotspot may confer an advantage in an evolving population, it is unclear how evolved motile isolates compare in swimming competition assays. In this study, we selected the fastest individual non-hotspot mutants from race assays (Fig. 3) and competed them against the common hotspot mutant (*ntrB* A289C) in equal proportions on LB and M9 0.25% agar plates. One of the fastest Pf0-2x lines evolved in LB is *ntrB* Δ406-417 (in-frame ΔLVRG amino acid deletion). The fastest M9 evolved Pf0-2x mutant is *ntrB* G421C (A141P amino acid substitution).

The hotspot mutant was genetically engineered with a kanamycin resistance cassette, enabling differentiation of lines post-competition (see Methods). Growth curve phenotyping assays were performed to ensure there were no metabolic differences between the hotspot and non-hotspot lines (LB *p* =0.4373, M9 *p*=0.4373); Fig.S.5)). Competition assays between evolved motile isolates were performed. Motility agar plates (*n*=6 for each competition) were inoculated with equal numbers of cells of each strain (Pf0-2x-sm *ntrB* A289C *glmS*::Kan^R^and either Pf0-2x *ntrB* Δ406-417 or Pf0-2x *ntrB* G421C). The plates were incubated until the bacteria saturated the motility agar (>7 days after inoculation). The whole agar plate of each replicate was disrupted, serially diluted and spread onto selective agar to determine the relative abundance of the hotspot and non-hotspot mutants.

In an M9 environment, the M9-adapted non-hotspot mutant *ntrB* G421C consistently outcompetes the hotspot mutant (*p*=0.0039, Kruskal Wallis Rank Sum Test). Even in LB (its non-adapted environment), *ntrB* G421C constitutes a significantly higher proportion of the population in competition assays (*p*=0.008, Kruskal Wallis Rank Sum Test) although both are at detectable levels (Fig. 4). In contrast, the fastest LB-adapted non-hotspot mutant *ntrB* Δ406-417 persists in both the LB and M9 nutrient environment against the hotspot mutant, but interestingly, the hotspot mutant outcompetes *ntrB* Δ406-417 in its adapted environment, LB (*p*=0.03, Kruskal Wallis Rank Sum Test). Yet, *ntrB* Δ406-417 showed comparable persistence to the hotspot mutant in its non-adapted M9 environment (*p*=0.26, Kruskal Wallis Rank Sum Test). Growth differences may account for these observations. In shaking broth, the *ntrB* G421C mutant grows equivalently to the hotspot mutant in LB, but much better than the hotspot in M9. The *ntrB* Δ406-417 mutant grows equivalently to the hotspot mutant in LB and M9 (Fig.S.7). In competitive growth assays in shaking broth, we see very similar results (Fig. S.8). These results suggest that the fastest evolved non-hotspot mutants can persist and outcompete the common hotspot mutant in competitive swimming assays, even in environments that are not their adapted environment.

**Fig. 4.**
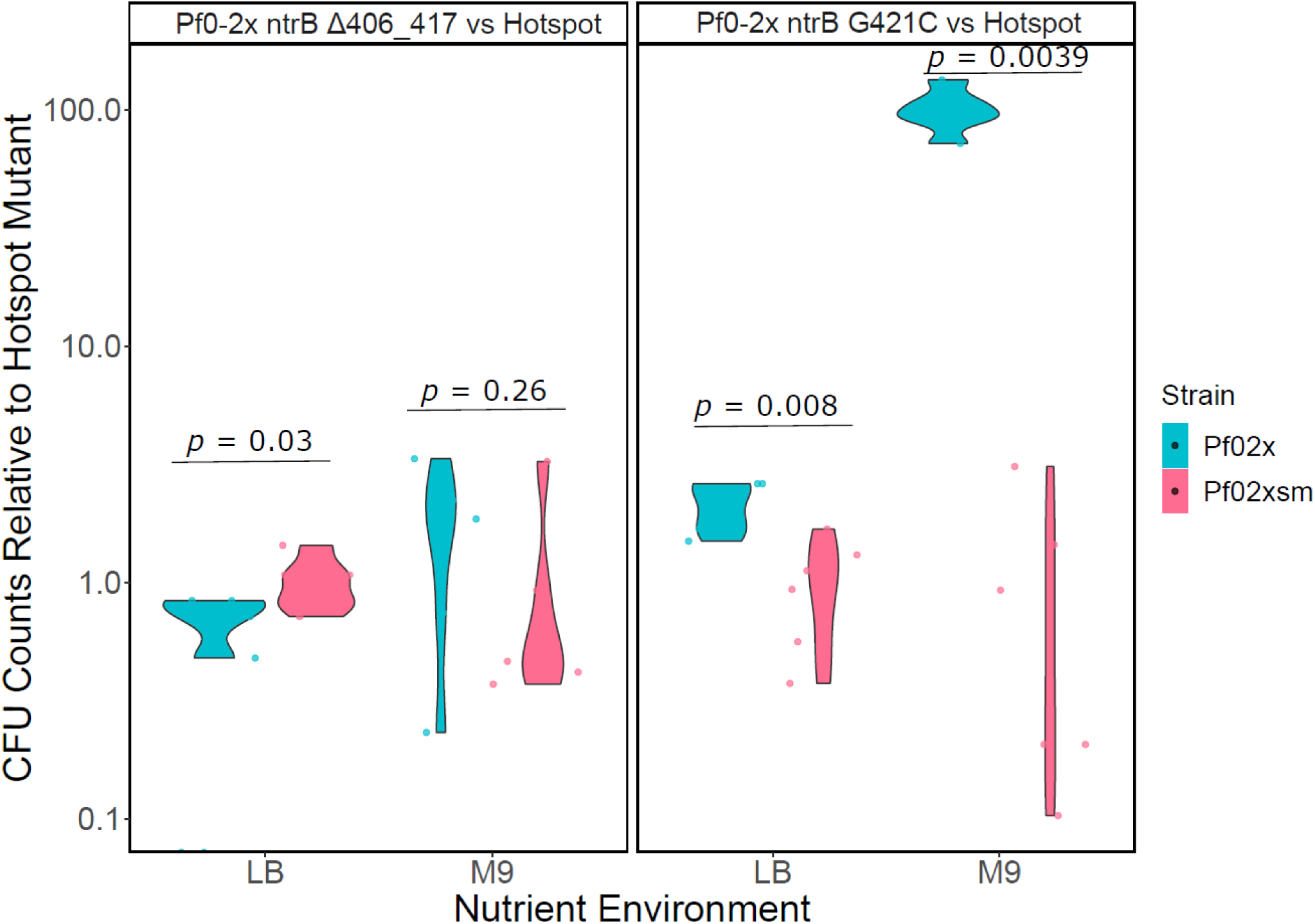
The persistence and abundance of the non-hotspot fastest swimmers in competition with the common hotspot mutant. The fastest non-hotspot (Pf0-2x) evolved lines (LB evolved Pf0-2x *ntrB* Δ406-417 and M9 evolved Pf0-2x *ntrB* G421C) were competed against the hotspot mutant (Pf0-2x-sm *ntrB* A289C *glmS*::Kan^R^) in 0.25% motility agar in both LB and M9 nutrient environments (*n*=6 for each competition). Colony forming units (CFUs) are given as a log value relative to the hotspot evolved strain.

## DISCUSSION

Mutational hotspots are facilitators of parallel evolution, especially when they work in concert with natural selection (1,38–40). But what role do they play in facilitating adaptation? Do they provide a competitive edge or an evolutionary handicap? Here, we look at the evolutionary rescue of the bacterial flagellar network in *P. fluorescens* Pf0-1 when the master regulator for the flagellar regulatory cascade (FleQ) has been disrupted and is rendered non-functional. Within this model system we compare bacteria with two different evolutionary potentials: one containing a mutational hotspot in *ntrB* (Pf0-2x-sm) that rapidly rescues flagellar driven motility via a highly parallel mutational route, and one without a mutational hotspot in *ntrB* (Pf0-2x), that rescues flagellar driven motility via a much broader mutational spectrum. In both cases rescue of the flagellar system is by gene regulatory network rewiring via the nitrogen regulation pathway, which has pleiotropic consequences for nitrogen assimilation. The hotspot and non-hotspot strains were put under selection to restore their lost swimming phenotype, and evolved motile mutants compared to see how the presence or absence of a strong mutational hotspot affected evolutionary trajectory and fitness. Under strong selection for motility in different nitrogen environments, we find that mutations in the hotspot strain are constrained to a single gene (*ntrB*) and almost exclusively to the same nucleotide (*ntrB* A289C). In contrast, in the non-hotspot strain we see a broader range of mutations that vary depending on the nutrient environment in which the bacteria were evolved.

We explored the influence of different nutrient environments on adaptive outcomes: LB (a complex medium containing a mix of carbon and nitrogen sources) and M9 (a defined nutrient medium containing a single nitrogen and carbon source). We saw a clear bias towards mutations in *glnA* when the non-hotspot strain was evolved in the complex nutrient media LB. On the other hand, we saw a higher frequency of mutations in *ntrB* when evolved in the defined nutrient media M9. This indicates that the nutrient environment Is having a direct influence on evolutionary outcomes in the non-hotspot strain (Pf0-2x), while the hotspot strain is constrained in both media environments. This may be as a result of differing metabolic stresses due to the difference in nutrient composition of each medium which may make some mutations in one setting less costly than in another (Fig.S.7). The influence of environment on evolutionary trajectory has already been shown elsewhere. Fraebel et al. (41) demonstrated that nutrient environment was the determining factor for adaptive strategy during multi-trait selection in *E*. *coli*. In complex media, faster migration was achieved through increased swimming speed and reduced growth, whereas in defined nutrient media this was achieved via amplified growth but reduced swimming speed.

Drawing on these findings, the nutrient environment not only influences the evolutionary trajectory and adaptive strategy, but it also appears to play a significant role in determining the mutational class that emerges when an organism responds to a particular selective pressure. In the evolved mutants derived from the non-hotspot strain, only SNPs were observed in the defined M9 environment, while in LB we observed a mix of SNPs and indels (*p*=0.001 Fisher’s Exact test). These results could be explained by changes in mutation bias introduced by environmental agents. Similar results have been reported in *E. coli* by Maharjan and Ferenci (42), where the influence of environment on mutational spectra was investigated through maintenance and evolution of replicate bacterial populations under chemostat-controlled nutrient limited conditions. Under complex nutrient conditions there was a high proportion of indels present, whereas in nitrogen limited conditions a higher proportion of SNPs were observed, similar to what we have seen in M9. We chose to look specifically at the impact of the nitrogen environment on mutational spectra due to the known pleiotropic costs of the motility restoring mutations on nitrogen assimilation. However, Maharjan and Ferenci (42) also noted mutational difference in other conditions limited in oxygen, iron, phosphorus and carbon. An alternative explanation for these results however is that, as the mutational spectrum changes between environments, more indels may be viable in LB or M9 than the other. This means it may present as mutation bias but instead be representative of the mutational spectrum. In either context these results highlight the power of environmental differences in making some mutational types more accessible than others. This in turn can have downstream effects on evolutionary outcomes and fitness as exemplified here, where mutations in different loci have distinct swimming and fitness profiles, with *glnA* mutants typically performing worse than *ntrB* and *glnK* mutants for the diverse mutant line Pf0-2x.

Our observations on the nutrient environment’s impact on the observed mutation spectrum, fitness, and swimming profiles of the evolved mutants serve as a suitable segue into our subsequent analysis. We proceeded to examine how the evolved swimming phenotype of these diverse evolved isolates, specifically those with distinct mutations, performed relative to the common hotspot mutation under different nutrient environments, and the implications this may have on the functional roles of the implicated genes. We found that the results were highly dependent on the nutrient environment in which the mutant line was evolved. Mutants evolved in M9 typically performed better relative to the common hotspot mutant, compared to those evolved in the LB environment. There were also a notable number of LB evolved mutants that would not swim in the alternate M9 nutrient environment (Fig.3). There were some reciprocal examples of this within the M9 evolved lines as well. Most of these were *glnA* mutants. GlnA is a glutamine synthetase enzyme, which aids in the assimilation of ammonia into the cell, and synthesises the amino acid glutamine (43,44). This enzyme is particularly important in Pseudomonads, as most cannot fix their own nitrogen (with the notable exception of *Pseudomonas stutzeri* (45)), and therefore rely on it heavily to provide the components for amino acid biosynthesis. While the specific effects of the mutations in *glnA* are unknown, the domains that they target are suggestive of a negative effect on the enzyme’s catalytic activity. One mutant had a stop codon following the second residue (*glnA* 2 ^Ser->STOP^, C5A), heavily suggesting the protein is non-functional as it is so severely truncated. In a recent study in *Salmonella enterica* Serovar Typhi, it was shown that deletion of *glnA* greatly increased expression of the NtrBC two component system (46). Taylor et al. (25) showed in their expression analyses of the model system used in this study that the means through which motility was initially restored was via hyper activation of NtrC. Effective deletion or inactivation of *glnA* would achieve this.

It seems likely that these *glnA* mutants were not able to swim in the alternate nutrient environment (M9) from which they had evolved because of *glnA*’s key role in ammonia assimilation. In M9 media the sole nitrogen source is ammonia so if a key protein to allow this into the cell is non-functional, the cells will likely be incapable of assimilating any nitrogen, therefore making it near impossible for them to carry out any basic cellular functions. Therefore, *glnA* mutations may be too costly to host in this environment, especially those that render it completely non-functional. In a complex media such as LB, it is possible for the cell to access nitrogen from multiple sources such as histidine (a carbon and nitrogen containing compound, which could use an alternative *cbrAB* two component system to assimilate nitrogen into the cell (47)), meaning mutations in *glnA* will not be as detrimental to the bacteria as they can assimilate nitrogen using alternative pathways. However, when grown free from selection for a swimming phenotype in shaking LB and M9 (Fig. S.7) we saw that there was no significant difference in the growth of the non-swimmer *glnA* mutants (*p*= 0.146), but this may also be due to reversion of bacteria to a non-flagellate form, as the benefit of maintaining the flagella is outweighed by the cost of impaired nitrogen regulation in shaking broth. Reversion to an immotile state was frequently observed throughout this research for mutants that seemed to have a very high metabolic cost.

Finally, we tested the rate of evolutionary rescue to restore the swimming phenotype between the hotspot and non-hotspot mutational lines in equally mixed populations in LB and M9 0.25% agar. We found that the hotspot dominated in both environments (LB: 6/6, M9: 5/6). This highlights the importance of mutational accessibility when in competition. As the hotspot strain was able to realise and fix the motility-restoring mutation first, it was able to dominate the ecological space. The importance of timing in establishment and survival in a niche is already well known in ecology, where “priority effects” (the sequence of arrival to an environment), determines the outcomes and interactions of groups within that community (8,48– 50). A similar phenomenon seems to be prevalent here with regard to mutational accessibility; initial establishment seems to be the most important factor for persistence rather than the fitness of the mutation.

There is a growing body of literature that shows that mutation bias can be an important driver of adaptive outcomes (15,51–53). Our results demonstrate that mutational biases can facilitate adaption, if they provide quick access to mutations conferring a trait under selection. However, we also show that having access to a broader mutational spectrum can allow the exploration of a wider range of mutational options that may confer fitter phenotypes (54). Broadly, whether rapid, repeatable mutation, or access to a broad mutational spectrum is the best facilitator of adaption will depend on: the rate at which the hotspot can confer a phenotype under selection relative to alternative mutational routes; and the fitness of the hotspot mutation relative to other possible routes. This model system, which utilises a “buildable” mutational hotspot, in combination with theory presents an exciting opportunity to expand our knowledge in this area. In addition, we show that the relative fitness of possible mutational routes is likely to be highly dependent on the environment, and by extension, the stochasticity of the environment. Being able to adapt rapidly might be advantageous, but not necessarily if the hotspot confers a sub-optimal phenotype (27) compared to other mutational routes that are masked by its effect. Understanding the role of mutation bias in defining adaptive outcomes remains an open and exciting question in evolutionary biology. Defining the nuances and details of when mutation bias can facilitate adaptive outcomes may present an opportunity to use this knowledge to improve our ability to forecast evolutionary trajectories of evolving microbial populations.

## AUTHOR CONTRIBUTIONS

LM Flanagan wrote the manuscript and collected the data for this publication (except where stated otherwise within the article). JS Horton and TB Taylor contributed to research conceptualisation and edits to the manuscript.

## CONFLICTS OF INTEREST

The authors confirm there are no conflicts of interest in this work.

## FUNDING INFORMATION

This project was funded by a Royal Society Enhancement Grant (RGF\EA\180265; awarded to T.B.T.) supporting L.M.F., BBSRC NI grant (BB/T012994/1; awarded to T.B.T.) supporting J.S.H. and a Royal Society Dorothy Hodgkin Research Fellowship (DH150169) awarded to and supporting T.B.T. Illumina WGS was performed by SeqCenter (LLC), Pittsburgh, PA, USA. Figures S1 and S2 were created using BioRender.com.

## ACKNOWLEDGEMENTS

The authors would like to thank the Royal Society for funding this research. They would also like to thank Matthew Shepherd for invaluable advice and guidance with the genetic engineering aspects of this project. They would also like to sincerely thank Kees Wanders for help with statistical analyses in this work.

## SUPPLEMENTARY MATERIALS

**Table S.1.** Strains used in this study.

| Strain | Specification | Growth Condition | Source |
| --- | --- | --- | --- |
| Pf0-2x | Immotile derivative of *Pseudomonas fluorescens* Pf0-1. Transposon insertion disruption of *fleQ*. | Ampicillin 100 µg/µl<br>or<br>Streptomycin 50/250 µg/µl | (55) |
| Pf0-2x-sm | Transposon insertion disruption of *fleQ*, and 6 synonymous base changes in *ntrB* generating a mutational hotspot. | Ampicillin 100 µg/µl<br>or<br>Streptomycin 50/250 µg/µl | (8) |
| Pf0-1 delta *fleQ* | Deletion of *fleQ*. Functionally equivalent to Pf0-2x. | - | Gifted by Mark Silby |
| Pf0-2x-sm *ntrB* A289C *glmS*::Kan<sup>R</sup> | Evolved motile version of Pf0-2x-sm with a kanamycin resistance cassette inserted into the *glmS* locus via mini Tn7 transposition | Streptomycin 50/250 µg/µl + Kanamycin 50 µg/µl | This study |

**Fig.S.1.**
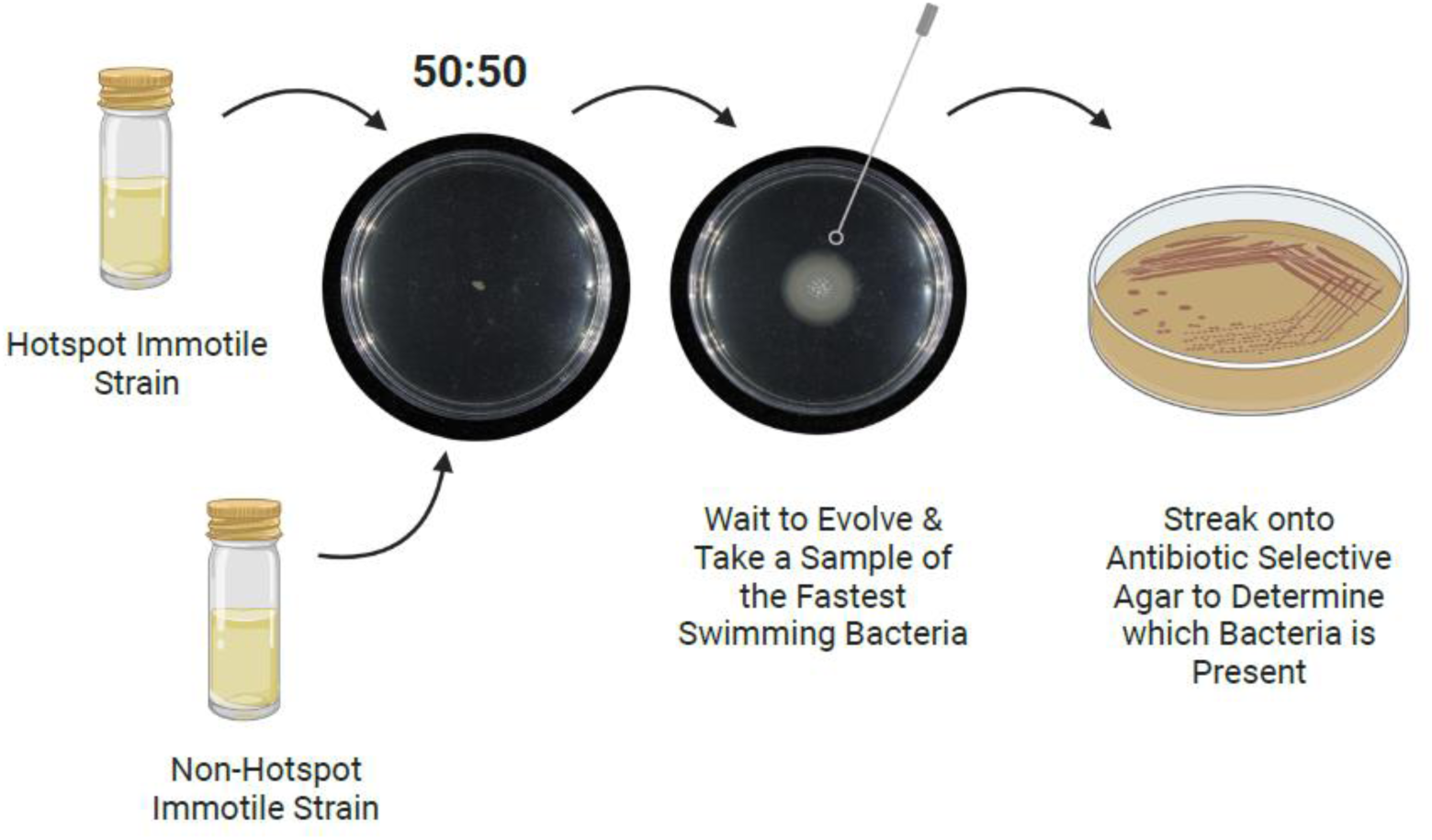
Experimental workflow of evolutionary rescue competitions. Created using BioRender.com

**Fig.S.2.**
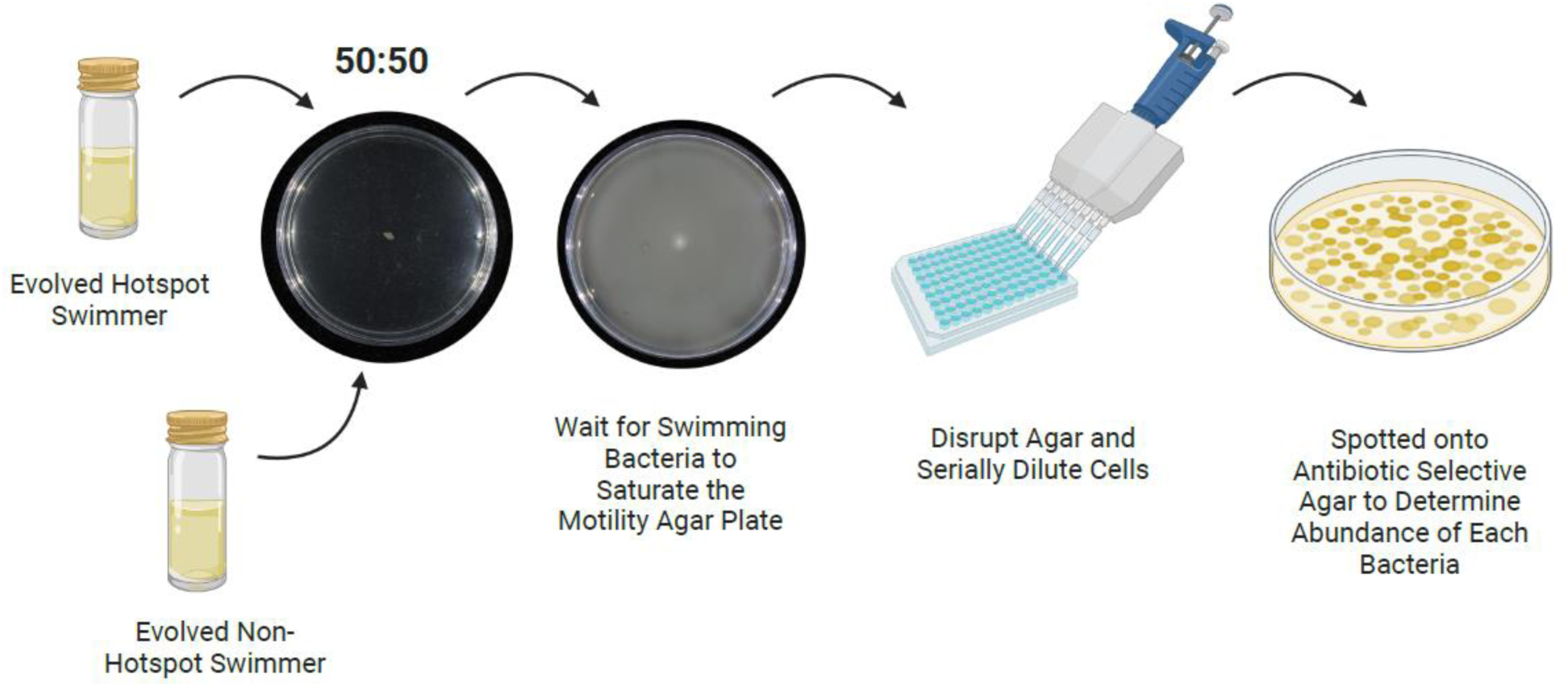
Experimental workflow of swimming competitions. Created using BioRender.com

**Fig.S.3.**
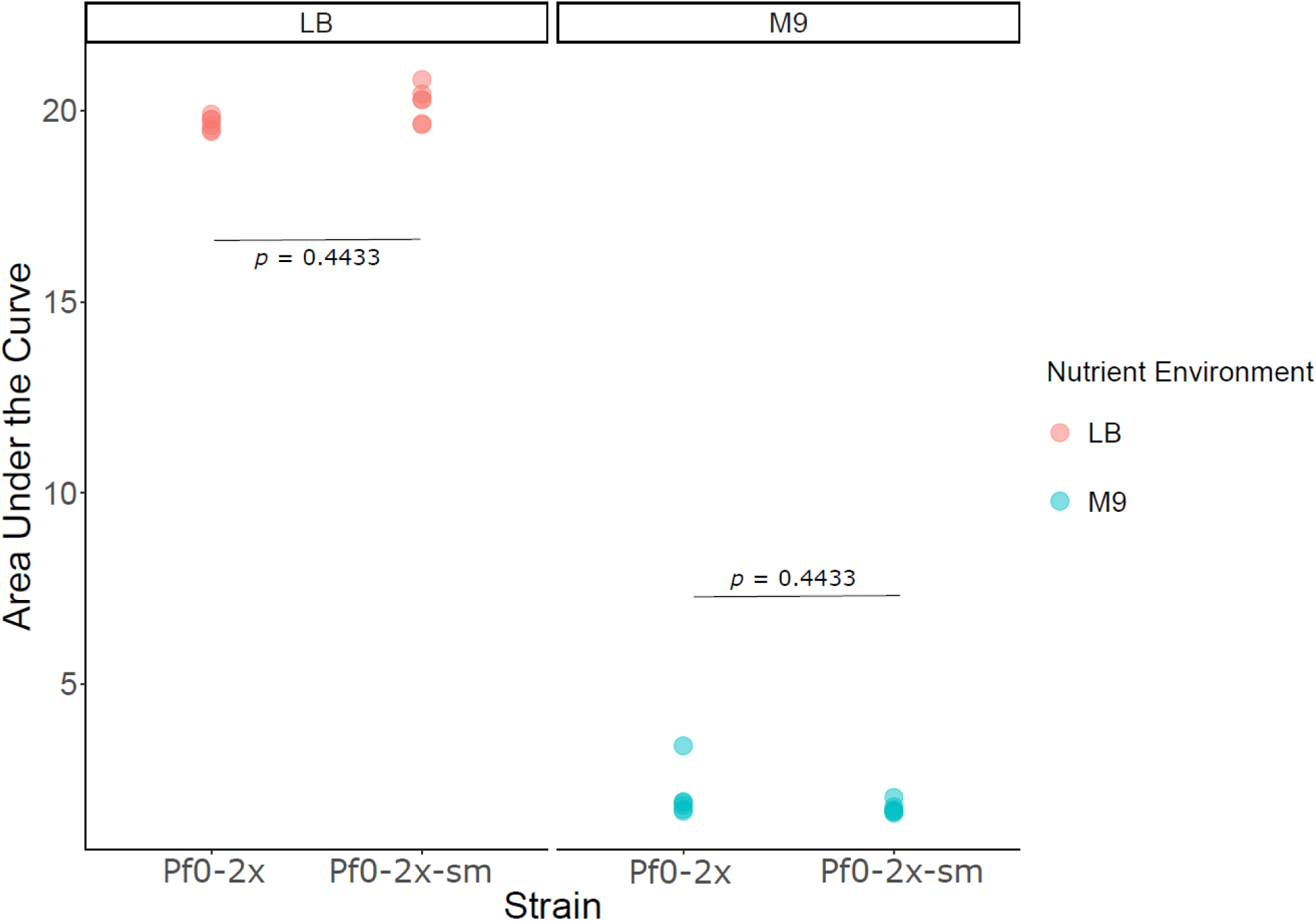
Measure of metabolic growth of Pf0-2x and Pf0-2x-sm.

**Fig.S.4.**
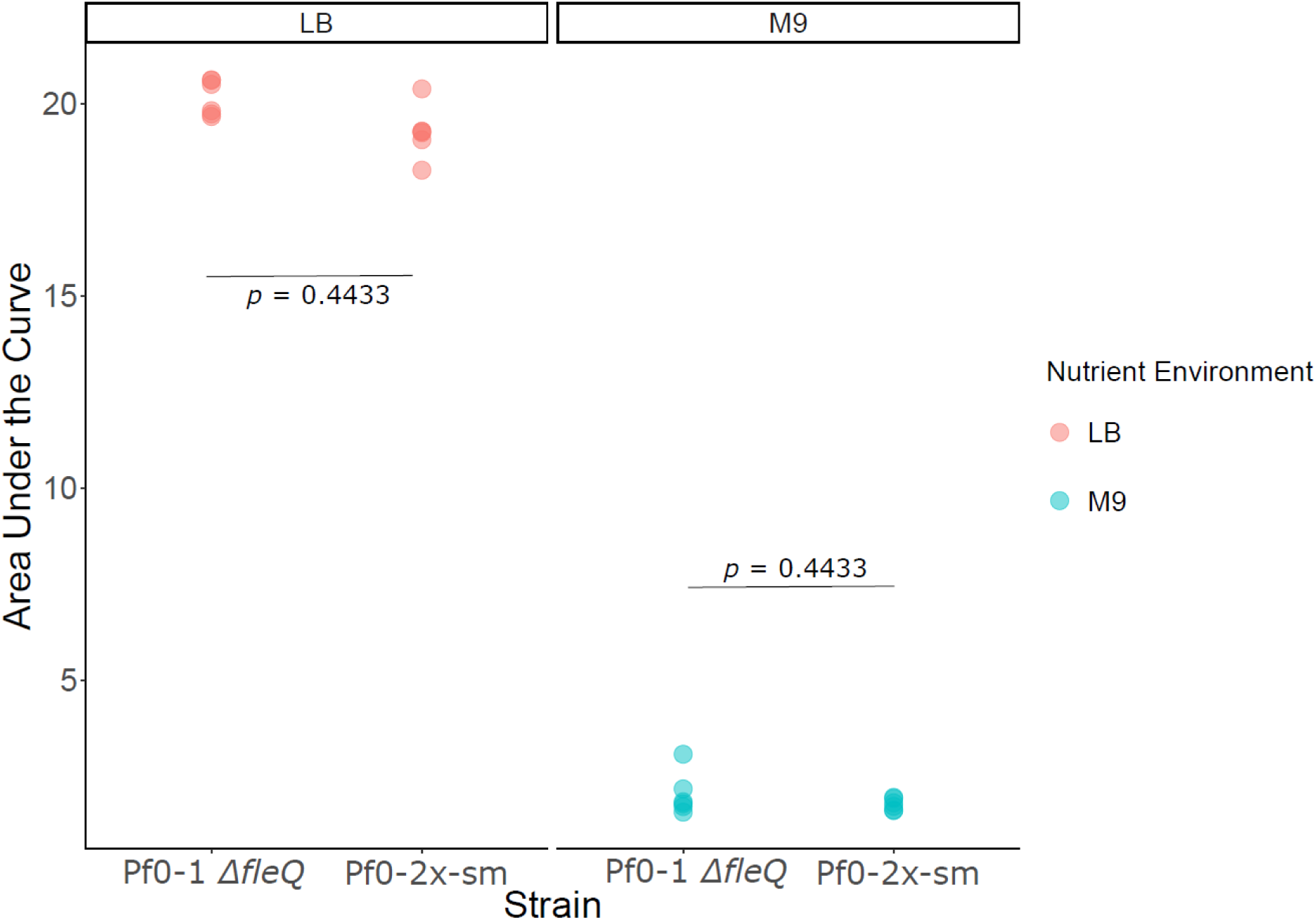
Measure of metabolic growth of Pf0-2x-sm and Pf0-1 Δ*fleQ*.

**Fig.S.5.**
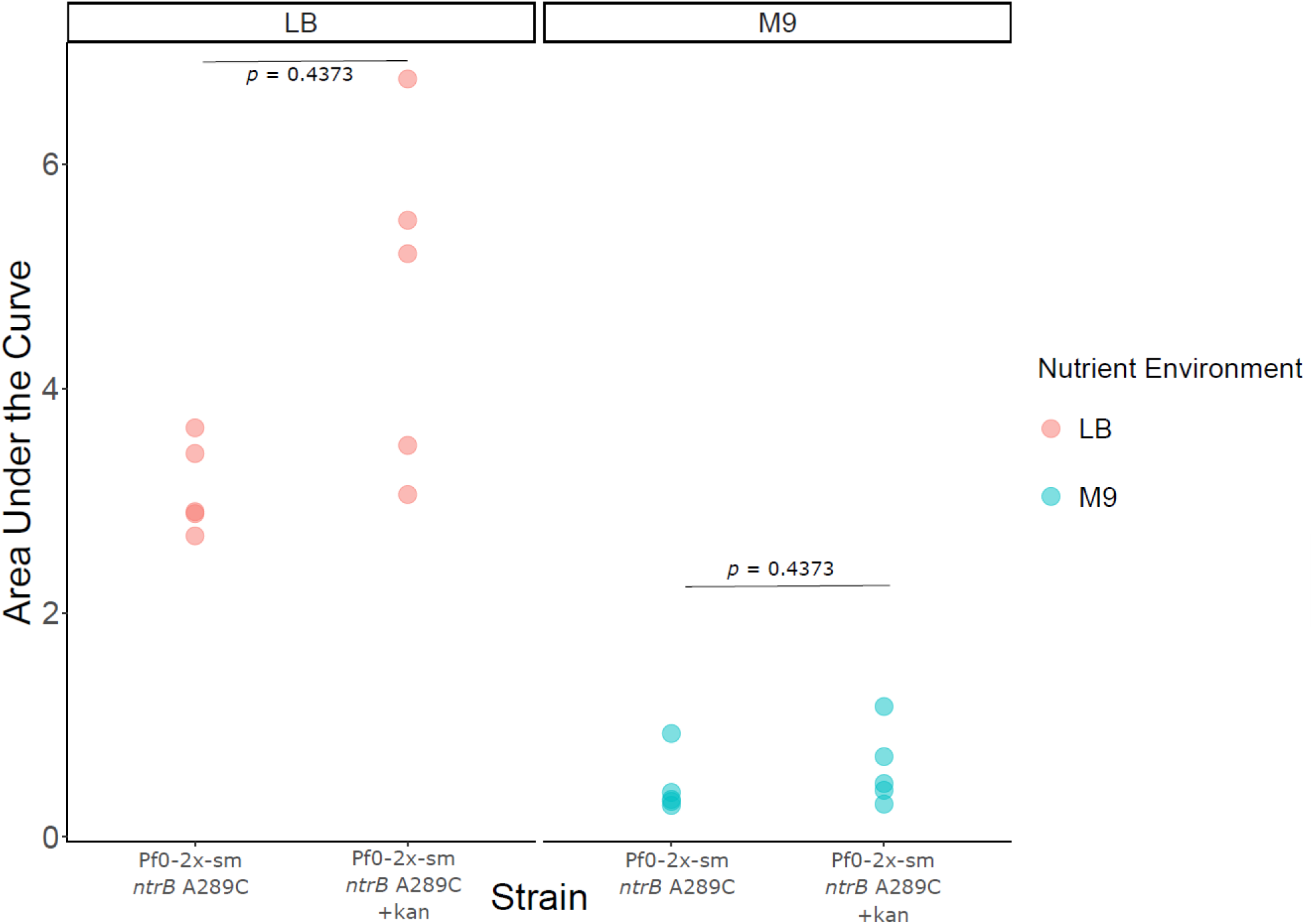
Measure of metabolic growth of Pf0-2x-sm *ntrB* A289C and Pf0-2x-sm *ntrB* A289C *glmS*::Kan^R^.

**Fig.S.6.**
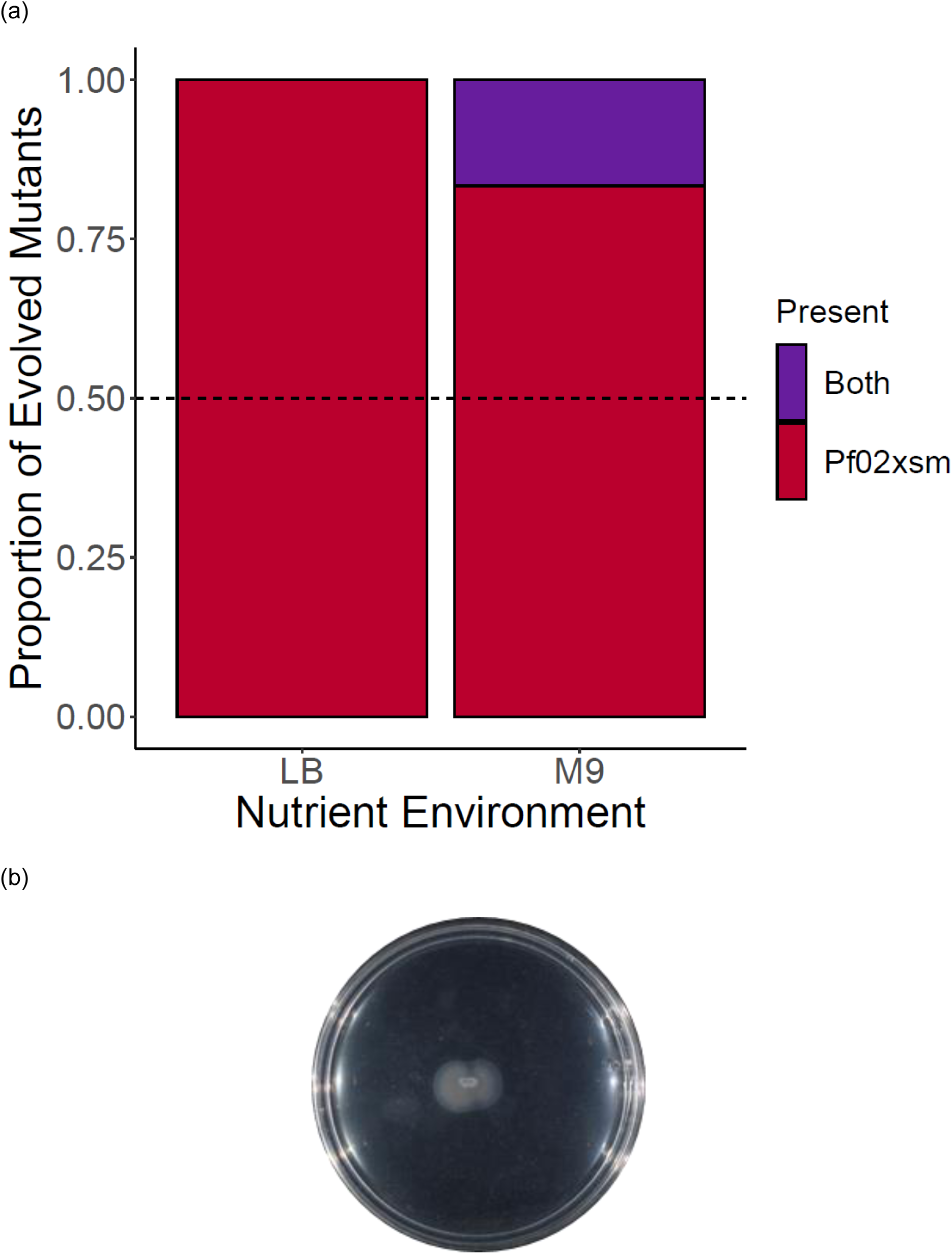
Competitive evolution of a motility phenotype between the immotile hotspot and non-hotspot lines in complex and minimal media. The hotspot (Pf0-2x-sm) and non-hotspot (Pf0-1 *ΔfleQ*) lines were inoculated in equal cell numbers (50:50) in LB (*n*=6) and M9 (*n*=6) 0.25% motility agar plates and motility allowed to evolve. When a motility phenotype was restored, the edge of the motile zone was sampled onto selective agar to determine which line had evolved first. **(a)** In the LB nutrient environment, the hotspot line was present at the leading edge in all replicates. In the minimal media environment M9, the hotspot line was present at the leading edge in 5 out of 6 replicates, but in one instance the non-hotspot line co-occurred at the leading edge of a second motile zone. The dashed black line crossing the y intercept at 0.5 indicates the expected values of each line if the competition outcome had been equal. **(b)** 0.25% M9 agar plate where the hotspot and non-hotspot lines evolved motility at the same time.

**Fig.S.7.**
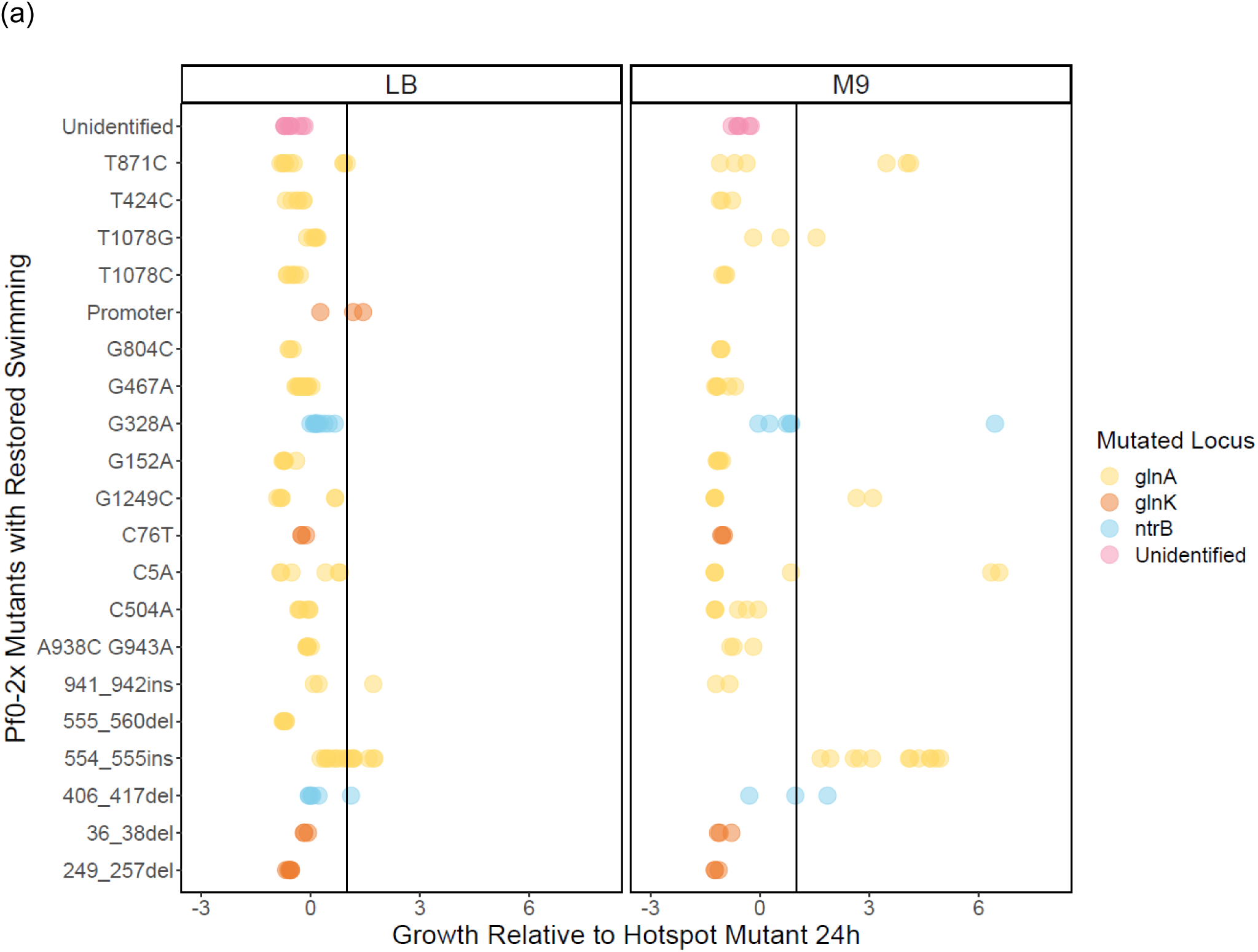
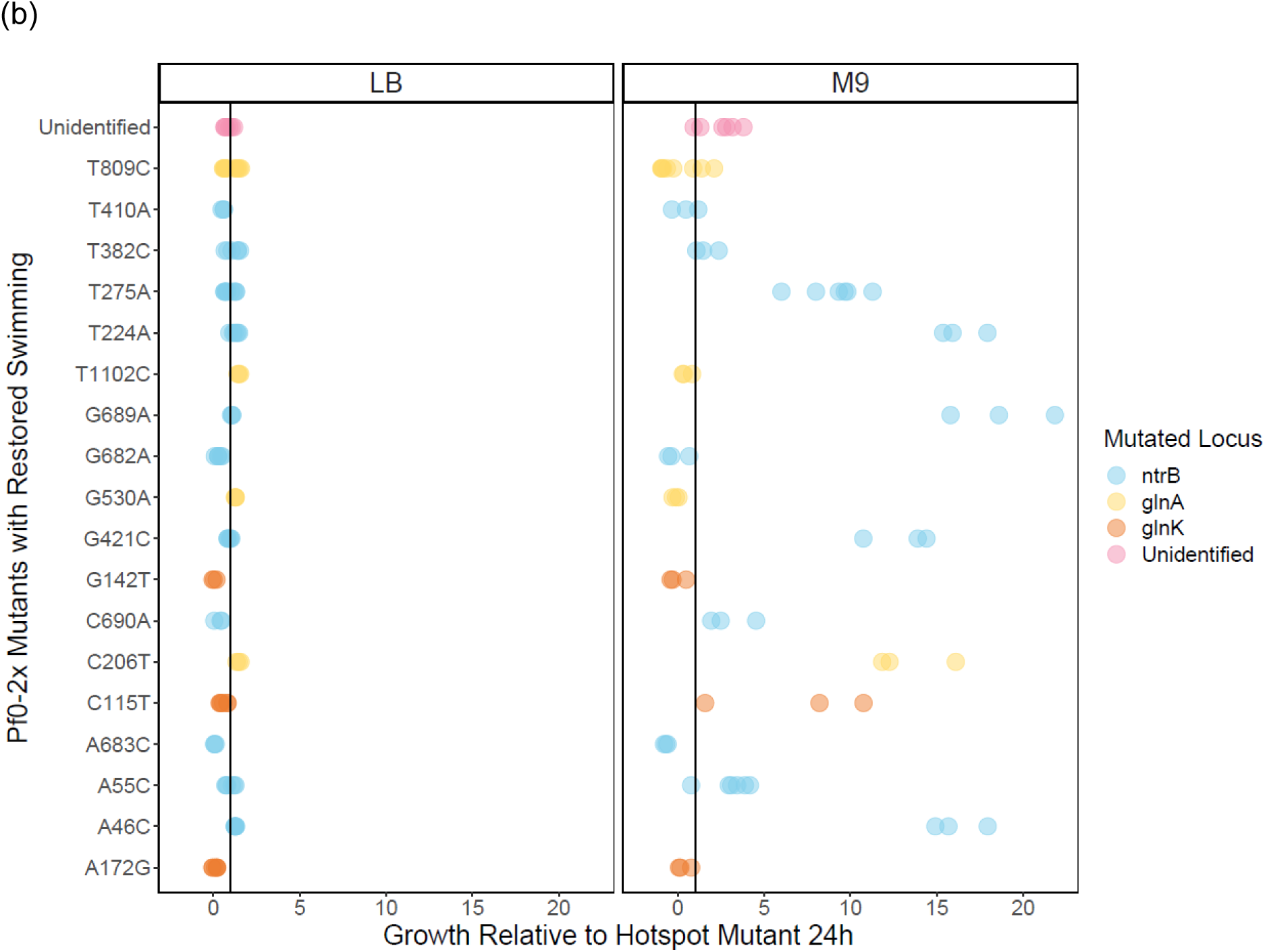
LB and M9 growth assay of all Pf0-2x evolved mutants relative to the common hotspot mutant. LB (a) and M9 (b) adapted Pf0-2x mutants (*n*=22, *n*=19) were assayed in LB and M9 in at least biological triplicate for 24h 180 rpm. The black line indicated the growth of the common hotspot mutant. Note that the x axis in the LB plot ranges from -3-8, while it ranges from -2-22 in the M9 plot.

**Fig.S.8.**
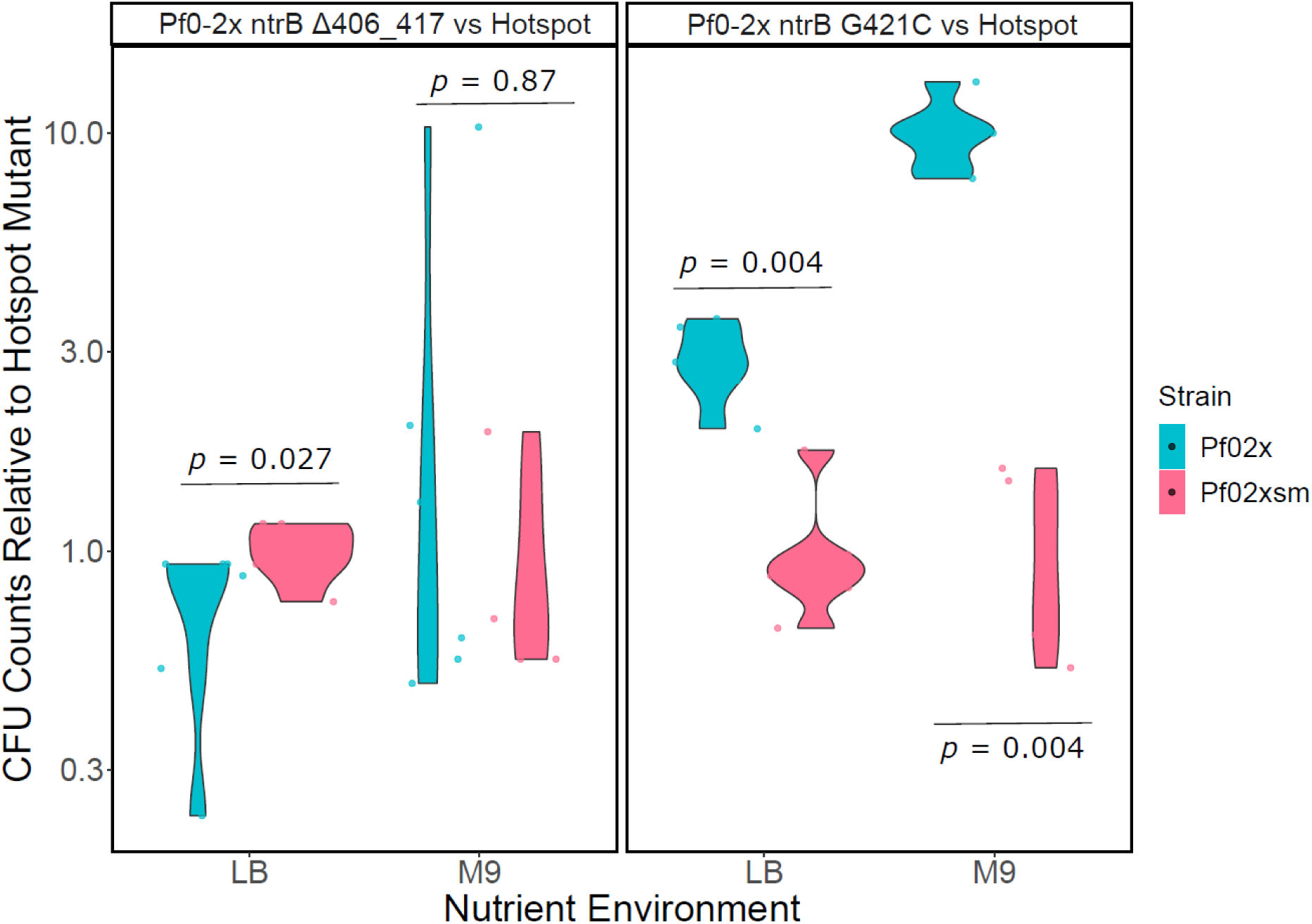
The relative abundance of the non-hotspot fastest swimmers in competition with the common hotspot mutant in shaking broth. The fastest non-hotspot (Pf0-2x) evolved lines (LB evolved Pf0-2x *ntrB* Δ406-417 and M9 evolved Pf0-2x *ntrB* G421C) were competed 50:50 against the hotspot mutant (Pf0-2x-sm *ntrB* A289C *glmS*::Kan^R^) in both shaking LB and M9 broth for 24 h (*n*=6 for each competition). Colony forming units (CFUs) are given as a log value relative to the hotspot evolved strain.

**Table S.2.**
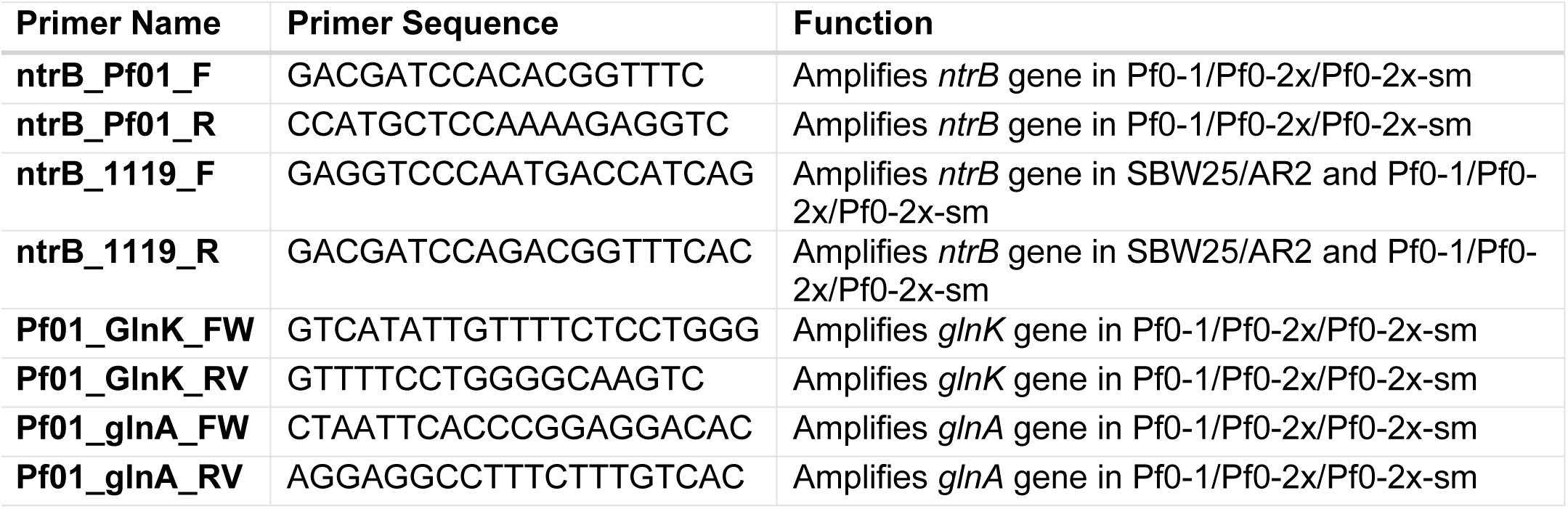
Primers used in this study.

## Notes

https://osf.io/8bt2w/?view_only=035fdd83de9f43bc9eeb70814a2ced46

### Competing Interest Statement

The authors have declared no competing interest.

